# Improved estimates for extinction probabilities and times to extinction for populations of tsetse (*Glossina* spp)

**DOI:** 10.1101/463067

**Authors:** Damian Kajunguri, Elisha B. Are, John W. Hargrove

## Abstract

A published study used a stochastic branching process to derive equations for the mean and variance of the probability of, and time to, extinction in population of tsetse flies (*Glossina* spp) as a function of adult and pupal mortality, and the probabilities that a female is inseminated by a fertile male. The original derivation was partially heuristic and provided no proofs for inductive results. We provide these proofs, together with a more compact way of reaching the same results. We also show that, while the published equations hold good for the case where tsetse produce male and female offspring in equal proportion, a different solution is required for the more general case where the probability (*β*) that an offspring is female lies anywhere in the interval (0, 1). We confirm previous results obtained for the special case where *β* = 0.5 and show that extinction probability is at a minimum for *β >* 0.5 by an amount that increases with increasing adult female mortality. Sensitivity analysis showed that the extinction probability was affected most by changes in adult female mortality, followed by the rate of production of pupae. Because females only produce a single offspring approximately every 10 days, imposing a death rate of greater then about 3.5% per day will ensure the eradication of any tsetse population. These mortality levels can be achieved for some species using insecticide-treated targets or cattle – providing thereby a simple, effective and cost-effective method of controlling and eradicating tsetse, and also human and animal trypanosomiasis. Our results are of further interest in the modern situation where increases in temperature are seeing the real possibility that tsetse will go extinct in some areas, without the need for intervention, but have an increased chance of surviving in other areas where they were previously unsustainable due to low temperatures.

**Author summary:** We derive equations for the mean and variance of the probability of, and time to, extinction in population of tsetse flies (*Glossina* spp), the vectors of trypanosomiasis in sub-Saharan Africa. In so doing we provide the complete proofs for all results, which were not provided in a previously published study. We also generalise the derivation to allow the probability that an offspring is female to lie anywhere in the interval (0, 1). The probability of extinction was most sensitive to changes in adult female mortality. The unusual tsetse life cycle, with very low reproductive rates means that populations can be eradicated as long as adult female mortality is raised to levels greater than about 3.5% per day. Simple bait methods of tsetse control, such as insecticide-treated targets and cattle, can therefore provide simple, affordable and effective means of eradicating tsetse populations. The results are of further interest in the modern situation where increases in temperature are seeing the real possibility that tsetse will go extinct in some areas, but have an increased chance of surviving in others where they were previously unsustainable due to low temperatures.

## Introduction

Whereas deterministic models of the growth of populations of tsetse fly(*Glossina* spp). (Diptera: Glossinidae) are adequate for large populations [1, 2], stochastic models are more appropriate when numbers are small, particularly if the population approaches zero through natural processes and/or following attempts to eradicate the fly. At that point the focus changes from attempting to attain deterministic predictions of future population levels, to predicting the probability that the population will go extinct, and the expected time required in order to achieve this end. Hargrove developed a stochastic model for the life history of tsetse flies (*Glossina* spp) and thereby provided estimates of the probability of extinction, and expected time to extinction, for these insects [3]. Such estimates were always of interest in situations where there was pressure in favour of area-wide eradication of entire tsetse species [4]. The model provided estimates of the level, and duration, of control effort required to achieve eradication of a target population and could thus be valuable for financial planning of tsetse and trypanosomiasis control efforts. The formulae developed were shown to provide good estimates of the time to extinction in successful operations that had already been carried out.

With the significant increases in temperature that have occurred over recent decades the model has assumed increased interest. It is becoming apparent that parts of Africa are becoming so hot that tsetse may no longer survive there. A well-documented example is the population of *G. pallidipes* Austen in parts of Zimbabwe. Whereas this species occurred in huge numbers in the area, for example, in the neighborhood of Rekomitjie Research Station, in the Zambezi Valley, the population has shrunk by *>* 99.99% over the past 30 years and now appears to be on the brink of disappearing [5, 6]. At the same time, other parts of Zimbabwe, where tsetse do not currently occur – in part because winter temperatures are too low – may soon be warm enough to support tsetse. Hwange National Park, for example, supported tsetse populations prior to the rinderpest epizootic of 1896: the fly never re-established itself in the area in the 20th Century, despite the presence of an abundance of wild hosts. In part this is due to the area always having been marginal climatically: increasing temperatures may change this balance in favour of the fly.

The above considerations prompted us to revisit the original derivations, from which several things became apparent: (i) It was assumed in the original derivation that equal proportions of male and female offspring were produced by female tsetse. The equations presented were correct for this particular case – but require modification for the more general case where the probability (*β*) that an offspring is female lies anywhere in the interval (0,1). (ii) At a number of points in the development it is claimed that results can be shown by induction, but the proofs are not provided. (iii) An heuristic explanation for one of the equation is misleading because it refers to a number *>* 1 as a probability. (iv) Finally, the development is restrictive in that it only treats the case where birth and death rates are constant over time. In the current paper we correct the first three problems and suggest ways of overcoming the fourth.

## Materials and methods

In this paper, we provide full details of the derivation of the formulae used and also provide a general form of the governing equation in [3], which can accommodate all possible values of *β*.

### Model Assumptions and Development

A female tsetse fly generally mates only once; it is thus crucial to include in our model the probability that a female tsetse fly is inseminated by a fertile male. We will also assume that the probability that a deposited pupa is male or female can be anywhere in the open interval (0, 1). Note that, at both endpoints, extinction occurs with probability 1.0, because the population will consist only of one gender of fly.

### Parameters and Interpretations

*λ* daily survival probability for adult female tsetse

*ψ* daily mortality rate for adult females = -*ln*(*λ*)

*φ* daily survival probability for female pupa

*χ* daily mortality rate for female pupae = -*ln*(*φ*)

*ν* time from adult female emergence to first ovulation(days)

*∊* probability female is inseminated by a fertile male

*τ* inter-larval period(days)

*P* pupal duration(days)

*p_n,k_* probability female tsetse fly dies between pregnancy *n* and (*n* + 1) and produces *k* surviving female offspring

*β* probability deposited pupa is female

The probability *p*_1,1_ that a female survives one pregnancy and produces one surviving female offspring is calculated as follows: First, we know that a female tsetse fly is inseminated by a fertile male with a probability *∊*, then survives with probability *λ*^(*ν*+*τ*)^ up to the time she produces her first pupa, which itself has a probability *β* of being female. This pupa survives the pupal period with a probability *φ^P^*, and the mother finally dies with a probability (1 – *λ^τ^*) during the next pregnancy. Thus, combining all these factors, we obtain the probability that a female tsetse fly produces one surviving daughter after surviving one pregnancy as

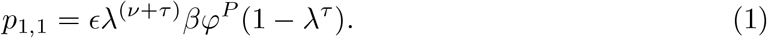

In general the probability that a female tsetse fly produces *k* surviving daughters after surviving *n* pregnancies is given by

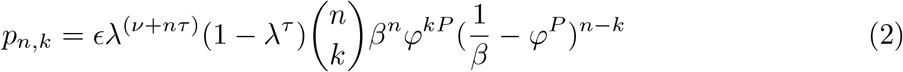

for *n >* 0, 1 *≤ k ≤ n*, and where (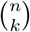) are the binomial coefficients.

#### Proof

Let *A_n_* be the event ‘a mother deposits exactly *n* pupae’, and *B_n,k_* be the event ‘*n* pupae produces exactly *k* female adults’. We can then define

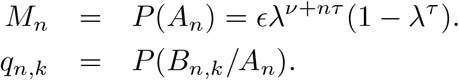

It is clear that

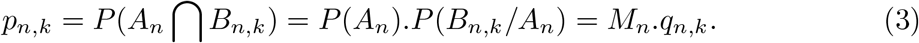

We notice that *M_n_* refers to the mother’s survival and *q_n,k_* refers to the pupae survival. So we can base our proof by concentrating on the pupal survival since the product of the two gives the result of interest.

It was actually observed that equation (2) can be proved without resorting to induction. Notice that for each pupa there are two possibilities; either it becomes an adult female or it does not. The probability that it becomes an adult female is *βφ^P^*, and the probability that it does not is then clearly (1 – *βϕ^P^*). Since the probabilities are the same for all pupae, and these outcomes for different pupae are independent, the probability that there are *k* adult females from *n* pupae is given by a binomial distribution as

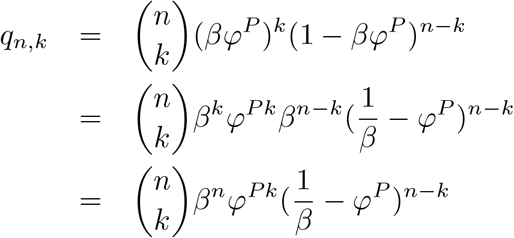

Thus, from equation (3), we obtain the expression for *p_n,k_* as

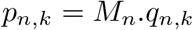

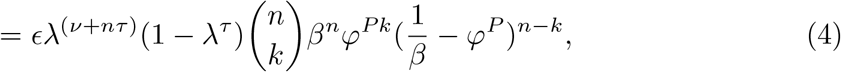

Note that this reduces to the governing equation in [3] when *β* = 0.5.

#### Remarks

1. The heuristic explanation for equation (2) in [3] is misleading because it terms a number greater than 1 a probability. Nonetheless, the formula is correct for the case he considered, and is also correct more generally with the adjustment of that term, as the proof shows.
2. The governing equation in [3] works only when *β*=0.5. After making the correction, it can be observed that equation (4) works for all values of *β*.

Summing equation (2) over *n* leads to the probability (*p_k_*) that a female tsetse fly produces *k* surviving female offspring before she dies. Thus

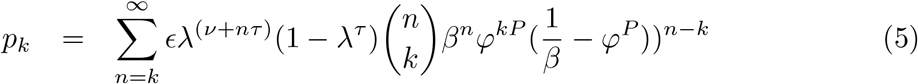

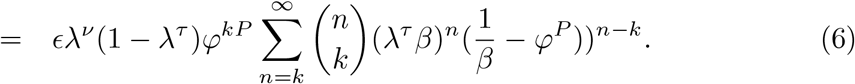

Thus, in general

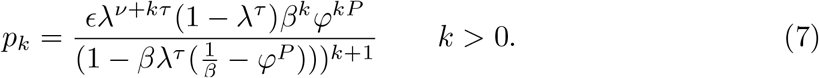

The probability that a female tsetse fly produces at least one surviving daughter before she dies can be obtained by summing equation (7) over *k >* 0, to obtain

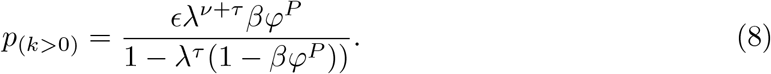

(See Supporting information for detailed proofs of equations (7) and (8))

Thus, the probability that a female tsetse fly does not produce any surviving female offspring before she dies is given by

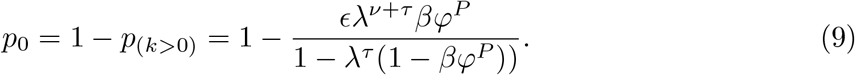

Assuming that we start with one female tsetse fly in the initial generation, which produces *k* surviving offspring, we can write the moment generating function for the next generation as

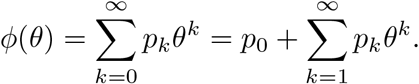

Substituting for *p*_0_ and *p_k_* and putting the terms not involving *k* outside the summation sign we get

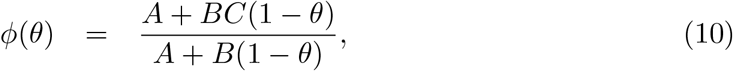

where *A* = 1 *− λ^τ^, B* = *βλ^τ^ φ^P^* and *C* = 1 *− ∊ λ^ν^*

The extinction probability can be found by solving the quadratic equation *ϕ*(*θ*) = *θ*, and it will be the smallest non-negative root [7, 8]. Thus the extinction probability is:

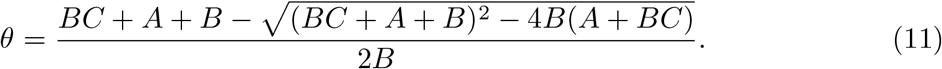

This is the probability that a female tsetse population, resulting from an initial population of one fly, goes to extinction. If the initial population consists of *N* flies, then, assuming the independence of the probability of extinction of each female line, the probability of extinction is *θ^N^*.

### Mean and variance of female tsetse population at generation n

We will use the method of moments to find the mean and variance of the expected number of offspring produced. From these variables we can then derive the mean and variance of the female tsetse population at a given generation *n*.

By definition, the *m*^th^ moment of *p_k_* is given by

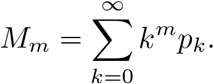

When *m* = 1, we obtain the first moment as

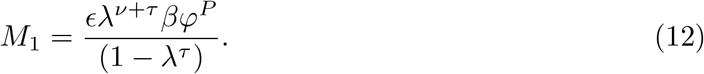

And when *m* = 2, we obtain the second moment as

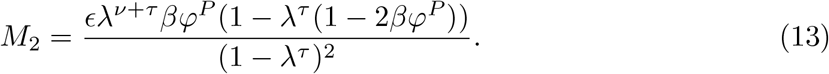

(See Supporting information for the proofs of equation (12) and equation (13))

The mean, or expected number of surviving daughters of female tsetse fly is

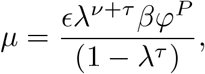

and the variance is given by

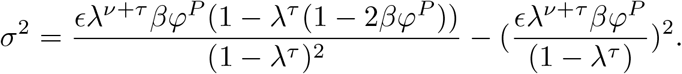

Where

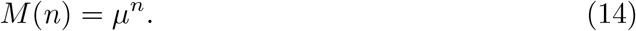

and

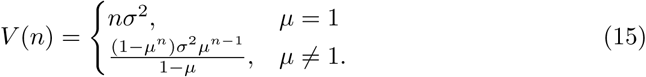

*M* (*n*) and *V* (*n*) are the mean and variance of the size of each generation (*X_n_*) respectively with the assumption *X*_0_ = 1. Equations (14) and 
(15) can be shown easily by induction.

### Time for population of the female tsetse flies to become extinct

From the general framework developed by Lange [7, 8] for the probability of extinction of a branching process. We have

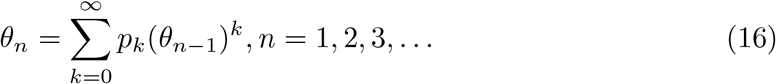

Where *θ_n_* is the probability of extinction at the *n^th^* generation and *k* is the number of offspring. Equation (16) can be rewritten in terms of a moment generating function as

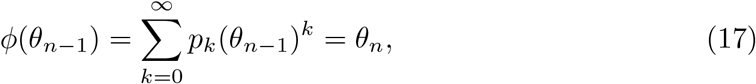

Thus, from (17), extinction probabilities can be calculated by starting with *θ*_0_ = 0*, θ*_1_ = *ϕ*(*θ*_0_)*, θ*_2_ = *ϕ*(*θ*_1_), and continuing iteratively through the generations to obtain

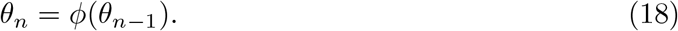

We also derived the first moments of *T*, based on the general formula obtained by Feller in [9] as

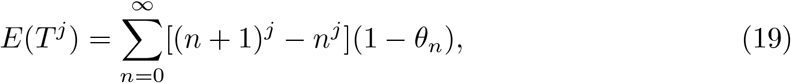

where (1-*θ_n_*) = P(*T > n*) and *T* is the extinction time. The first two moments of *T* are:

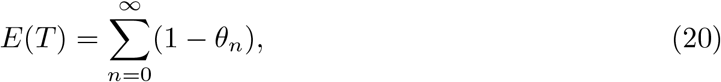
and

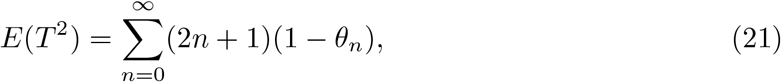

Thus, using equations (10) and (18) and taking *θ*_0_ = 0, we can calculate the values of *θ_n_* by iteration. The first two, for example, are:

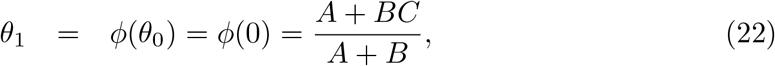

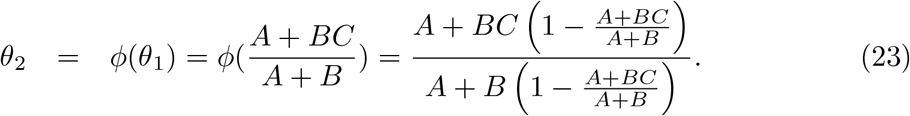

In a situation where there are *N* surviving females, with *N >* 1, equations (20) and (21) can be generalised. The probability of extinction at or before generation *n* is *θ_n_*. If we have *N* surviving females, then the probability that they all become extinct at generation *n* is (*θ_n_*)^*N*^. Thus,

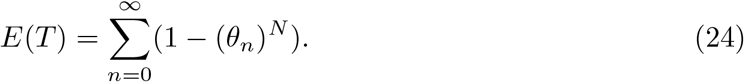

and

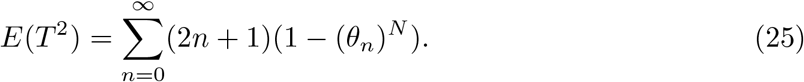

To estimate the mean and variance of the time to extinction for a population of *N* female tsetse flies, all that needs to be done is to estimate *θ_n_* for a population consisting of a single fly, raise each of the values to power *N*, and obtain the appropriate sums.

## Results

### Extinction probabilities as a function of adult and pupal female mortality rates

We produced MATLAB code to solve Equation 8 and generate the extinction probabilities for given values of parameters A, B and C. Our results were closely similar to those previously published [3], as illustrated in the Figures S1 - S6 in the Supporting information. For example, for a pupal duration (*P*) of 27 days, a time to first ovulation (*ν*) of 7 days, an inter-larval period (*τ*) of 9 days, a probability of *β* = 0.5 that a deposited pupa will be female and where all females are inseminated by a fertile male (*∊* = 1), the extinction probability for a population consisting of a single inseminated female fly increased linearly with adult female mortality rate (*ψ*), at a rate which increased with increasing pupal mortality rate (*χ*) (Fig S1).

If the pupal mortality is high enough, then the probability of extinction is high even if the adult mortality is low. For example if *χ* = 0.03 per day, then there is a greater than 40% chance that extinction will happen, even if the adult mortality rate is only 0.01 per day. Even when there was zero pupal mortality, however, extinction was certain when adult mortality rate approached levels of 0.04 per day. When the pioneer population consisted of more than a single inseminated female, the extinction probability was of course generally lower (Fig S2). If the pupal mortality rate was even 0.005 per day, however, all populations eventually went extinct, with probability 1, as long as adult mortality rate exceeded about 0.032% per day.

### Extinction probabilities as a function of the probability of insemination

In situations where, for example, sterile male tsetse are released into a wild population or where a population is extremely low, females may fail to mate with a fertile male and *∊* will then fall below 1.0. When the starting population was a single inseminated female, and with other input parameters as defined above, the extinction probability increased approximately linearly with increasing values of *∊* (Fig S3). Increasing the assumed value of the adult mortality rate (*ψ*) simply shifted the whole graph of extinction probability towards a value of 1.0, without changing the rate of increase of extinction probability with *∊*.

When the pioneer population was greater than 1, the relationship with *∊* was no longer linear (Fig S4) and, even when the starting population was only 16 inseminated females, the extinction probability was still effectively zero when the probability of fertile insemination fell to 50%. No population could avoid extinction, however, when *∊* was less than about 10%

### Extinction probabilities as a function of the probability a deposited pupa is female, and the death rate of adult females

Extinction is of course certain if a population consists only of one sex, but the probability of extinction goes to 1.0 more rapidly as the probability (*β*), that a deposited pupa is female, goes to 0 (all male population) than as it goes to 1 (all female population, Fig 1). For adult female mortality rates very close to zero, the extinction probability goes to 1.0 as *β* goes to zero: but, for higher adult death rates the limit is reached for values of *β >* 0. For example, when *λ* = 0.98, extinction is already certain once the female proportion among pupae drops to 30%. The minimum extinction probability always occurs for a value of *β >* 0.5, by an amount that increases as adult female mortality increases.

**Fig 1.**
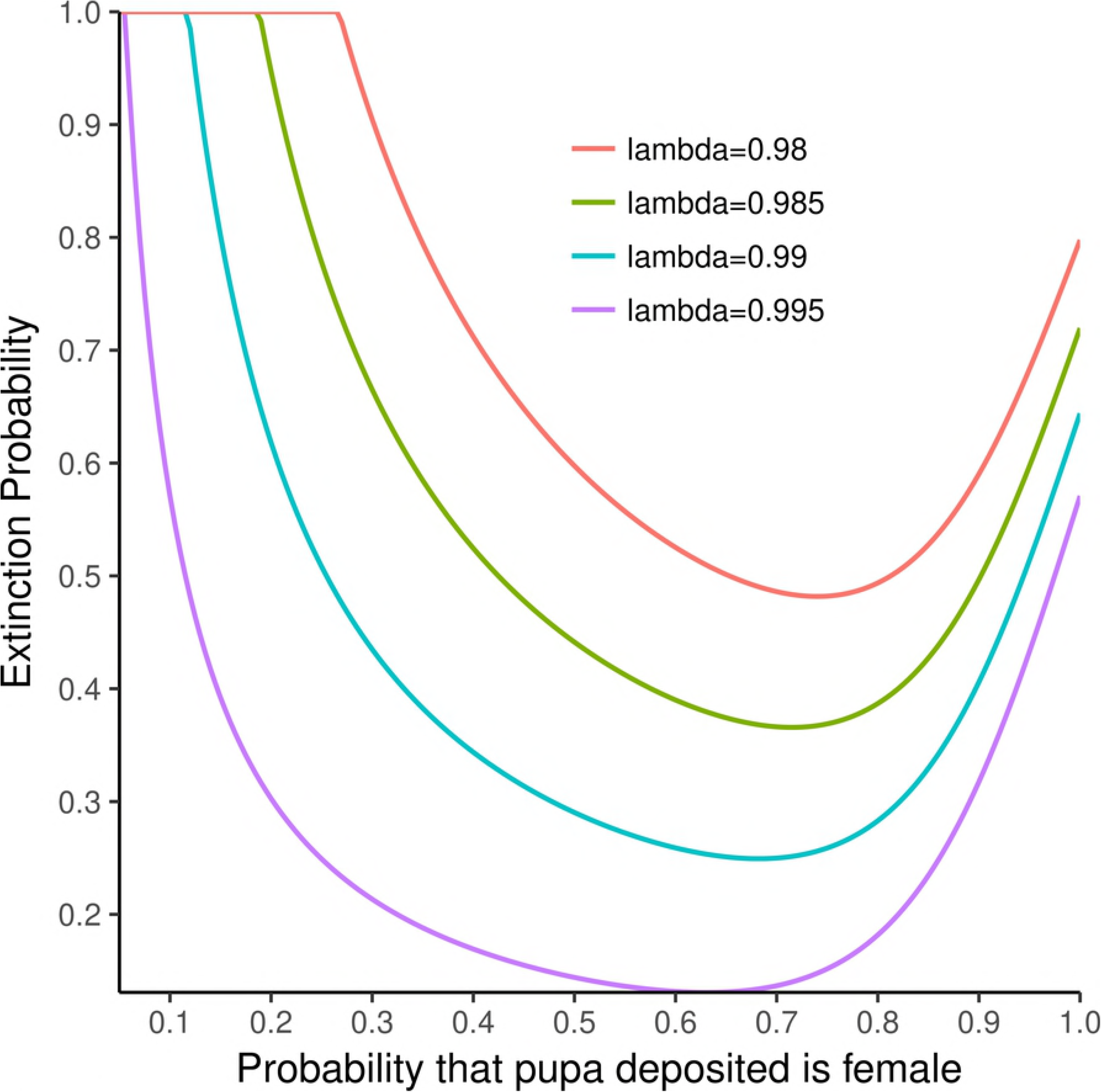
Extinction probability as a function of the probability a deposited pupa is female, and the adult survival probability. Input assumptions: Pioneer population *N* = 1 inseminated female; pupal mortality rate *χ* = 0.005 per day; probability female inseminated by a fertile male, *∊* = 1.0; pupal duration, *P* = 27 days; time to first ovulation, *ν* = 7 days; inter-larval period *τ* = 9 days. Figures in the body of the plot show the assumed daily survival probability (*λ*) for adult females.

### Expected number of generations to extinction

We derived the general equation for the expected number of generations to extinction for independent lines of *N* females in equation (21). Equations (19) and (20) give the first two iterations of the probability of extinction. MATLAB code was written to solve equation (21) iteratively and thus find the expected number of generations to extinction. Fig S5 shows that the expected number of generations to extinction decreases with any increase in pupal mortality.

Fig S6 gives the result of the expected number of generations to extinction against the probability of insemination. From the graph, we can see that the lower the probability of insemination by a fertile male, the smaller the number of generations to extinction.

Fig S7 shows that, in the event that eradication is attempted through the release of sterile males, in order to reduce the probability that females are inseminated by fertile males, the eradication process will be much hastened if the mortality of the wild female population is also increased.

## Discussion

Our results place on a firmer footing published findings based on the restrictive assumption that a deposited pupa has an equal chance of being male or female [3]. Nonetheless, we confirm various findings of the earlier study. For example, it is clear that tsetse populations can exist at very low population densities, and the sensitivity analysis added in the present study, also indicates the prime importance of mortality among adult females in affecting the probability of extinction in tsetse populations. This result further supports the arguments adduced in the earlier paper regarding the efficacy, and the cost-efficacy, of “bait methods” of tsetse control – and we refer the reader to the earlier discussion [3].

For controlling the species of tsetse occurring in Zimbabwe – *G. m. morsitans* and *G. pallidipes* – the primacy of bait methods has been well established. In a study carried out on Antelope Island, Lake Kariba, it was estimated that 24 odour-baited insecticide-treated targets, deployed on the 5 sq km island, killed about 2% per day of female *G. m. morsitans* and 8% of *G. pallidipes* [10, 11]. When targets were used in the Rifa Triangle, in the Zambezi Valley, at the same approximate density, populations of both species were reduced by *>* 99.99% and the populations in the treated area only survived through invasion from adjacent untreated areas [12, 13]. Again, it was estimated that the targets were killing about 2% per day of female *G.m. morsitans* and up to 10% of *G. pallidipes*. The Rifa experiment was carried out, however, when the only odour attractants available for use with targets were acetone and 1-octen-3-ol: targets are now used with the addition of two phenols, which increase the target kill rates of *G.m. morsitans* by about 50%, and those of *G. pallidipes* by several fold [14, 15]. Moreover, the targets currently in use are nearly twice as effective as the prototypes used in the Rifa Triangle [16]: consequently it is estimated that odour-baited insecticide-treated targets, deployed at 4*/sq* km in Zimbabwe will kill at least 4% per day of female *G. m. morsitans* and about 10% of female *G. pallidipes*. As is clear from Fig S1,these levels of imposed mortality are sufficient to ensure eradication of any population of tsetse, even if the natural, adult and pupal mortality rates are zero. These theoretical predictions are borne out for *G. m. morsitans*, which was eradicated in the Umfurudzi Safari Area using targets at the above density [17].

In areas of Zimbabwe where there are cattle in tsetse areas, the use of insecticide-treated cattle provides an effective method that can be used in parallel with, or even instead of, insecticide-treated targets. The combined use of these two bait methods saw massive reductions in levels of animal trypanosomiasis in north-east Zimbabwe during the 1990s, to the point that in 1997, despite widespread monitoring of cattle, no case of animal trypanosomiasis was detected [17].

In Zimbabwe, therefore, the use of any other method in addition to odour-baited insecticide-treated target, and insecticide-treated cattle, appears to constitute a waste of resources. In particular the release of large numbers of laboratory-reared sterile male tsetse appears superfluous and unjustified. In this regard the very much larger effect on the probability of extinction resulting from quite modest increases in adult female mortality stands in strong contrast to the very large reduction in female fertility that must be effected in order to achieve eradication (cf Figs S2 and S3).

Since the publication of the original analysis of extinction probability for tsetse in [3], there has been increased interest in using insecticide-treated targets in control operations against riverine species of tsetse, such as *G. f. fuscipes* and *G. palpalis* [ [18]-[25]]. These species are not strongly attracted by host odour, and the kill rate per target is thus very much lower than for *G. pallidipes*: the riverine species can, however, be captured on much smaller targets [typically 25 × 25*cm*] than those required for use with savannah species [up to 2 × 1*m*]. It is thus economically feasible to deploy much larger numbers of these so-called “tiny targets” and use them to effect significant control of riverine tsetse species. In a trial in northern Uganda where tiny targets were deployed at 20 targets per linear km (giving an average density of 5.7 per sq km), it was possible to reduce the fly population by*>* 90%. It was noted that this reduction was more than sufficient to break the transmission cycle for Human African Trypanosomiasis in the area,

In the same study, experiments on islands in Lake Victoria, Kenya, suggested that tiny targets used at the above density were killing 6% of the female population per day. The suggestion is that a further increase in target density might result in the eradication of populations, without the need to use any ancillary methods to control tsetse or trypanosomiasis.

## Limitation of the study

All of the results presented here have been calculated on the assumption that, for each scenario, all rates associated with rates of mortality and reproduction are constant over time. In reality, in the field, temperatures change with time and, since tsetse are poikilotherms, all of the mortality rates and developmental rates associated with reproduction also change continuously with time. The calculation of extinction probabilities is greatly complicated where temperatures are changing with time, and consideration of such situations is beyond the scope of the current study. Extreme weather events, such as prolonged spells of very hot weather, as have been experienced in recent years in the Zambezi Valley of Zimbabwe, may push tsetse populations close to extinction. We are currently investigating the circumstances under which it is be possible to calculate extinction probabilities in such situations. Where we cannot obtain analytical solutions of the type derived here, when all model parameters are time-invariant, we will use simulation methods to investigate the problem.

## Supporting Information Legends

**SI 1**: Proof of equation (7)

**SI 2**: Proof of equations (8 and (10)

**SI 3**: Proof of equations (12) and (13)

## Supporting Information Legends for figures

**Fig S1.**
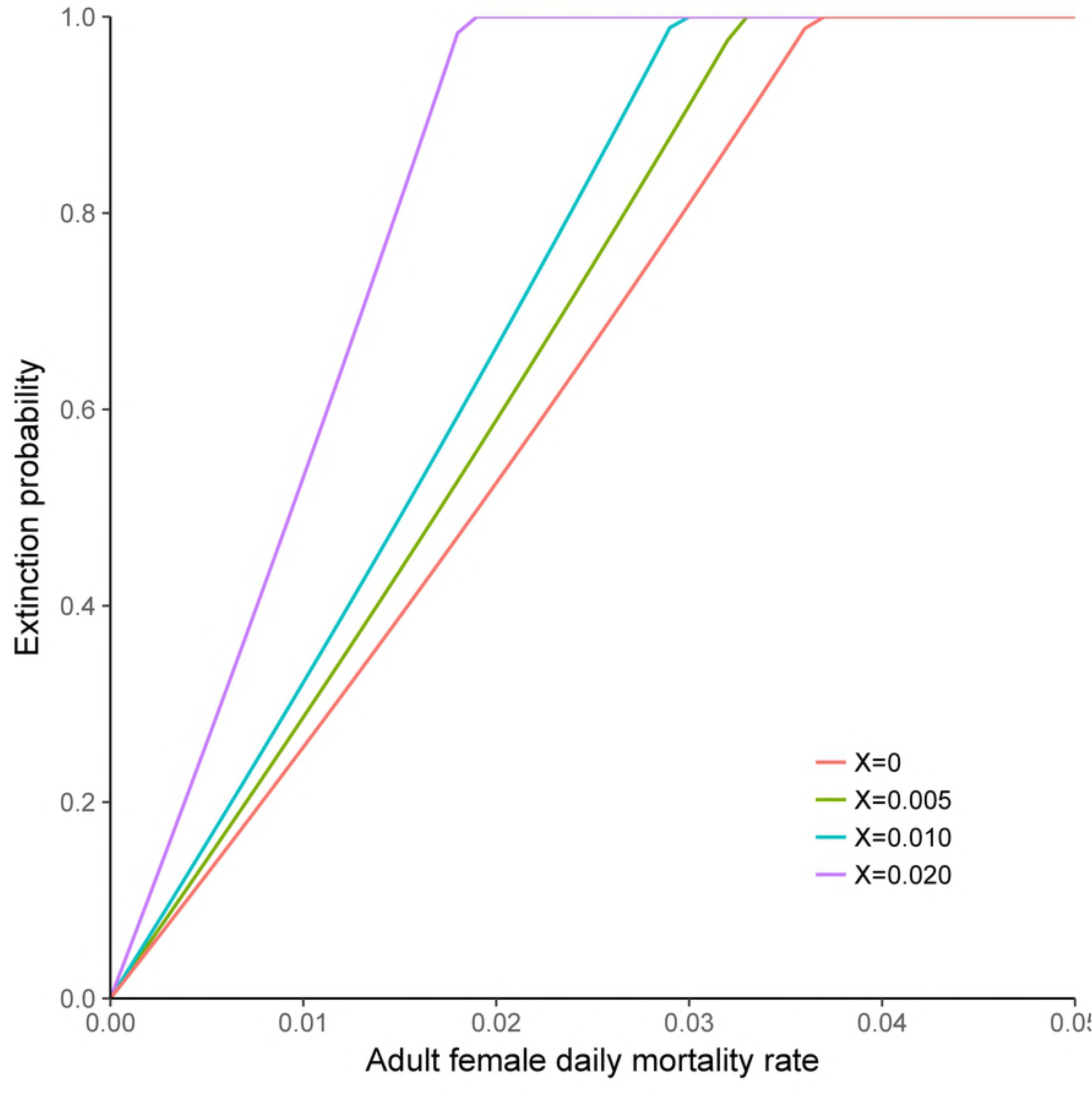
Extinction probability as a function of female adult, and pupal, mortality rates. Input assumptions: Pioneer population *N* = 1 inseminated female; probability females inseminated by a fertile male, *∊* = 1.0; probability deposited pupa is female, *β* = 0.5; pupal duration, *P* = 27 days; time to first ovulation, *ν* = 7 days; inter-larval period *τ* = 9 days. Figures in the body of the plot show the assumed pupal mortality rate (*χ* per day). (cf [3], Fig 1A)

**Fig S2.**
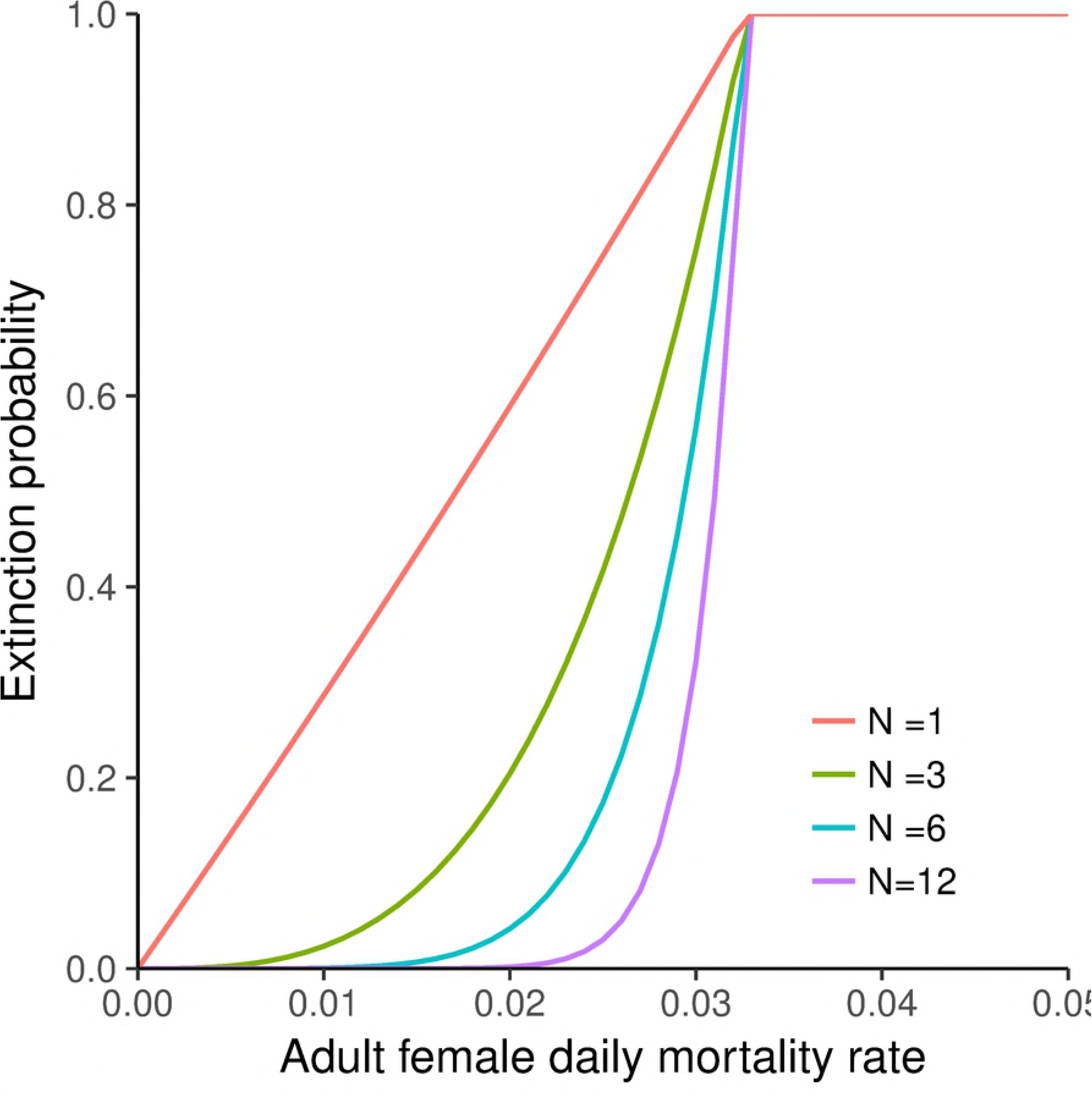
Extinction probability as a function of adult female mortality rate and the number of inseminated females in the pioneer population. Input assumptions: . Input assumptions: Probability females inseminated by a fertile male, *∊* = 1.0 probability deposited pupa is female, *β* = 0.5; pupal duration, *P* = 27 days; time to first ovulation, *ν* = 7 days; inter-larval period *τ* = 9 days. Pupal mortality rate assumed constant at a level of *χ* = 0.05 per day. Figures in the body of the plot show the assumed number (*N*) of inseminated females in the pioneer population. (cf [3], Fig 1B)

**Fig S3.**
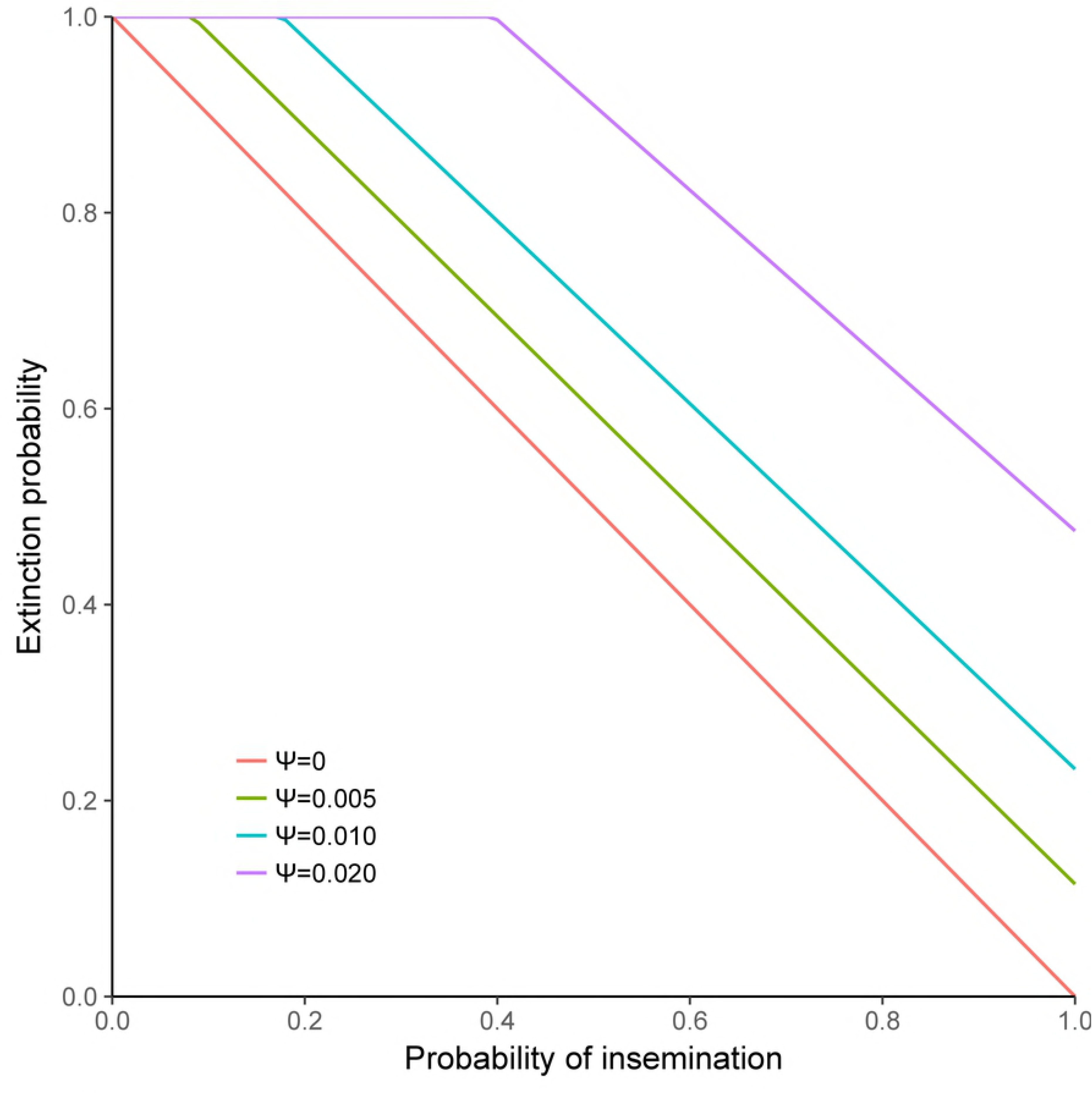
Extinction probability as a function of the probability that a female is inseminated by a fertile male, for different levels of adult female mortality rate. Input assumptions: Pioneer population *N* = 1 inseminated female; adult mortality rate *ψ* = 0.005 per day; pupal mortality rate *χ* = 0.005 per day; probability deposited pupa is female, *β* = 0.5; pupal duration, *P* = 27 days; time to first ovulation, *ν* = 7 days; inter-larval period *ν* = 9 days. Figures in the body of the plot show the assumed adult mortality rate (*χ* per day) (cf [3], Fig 2A)

**Fig S4.**
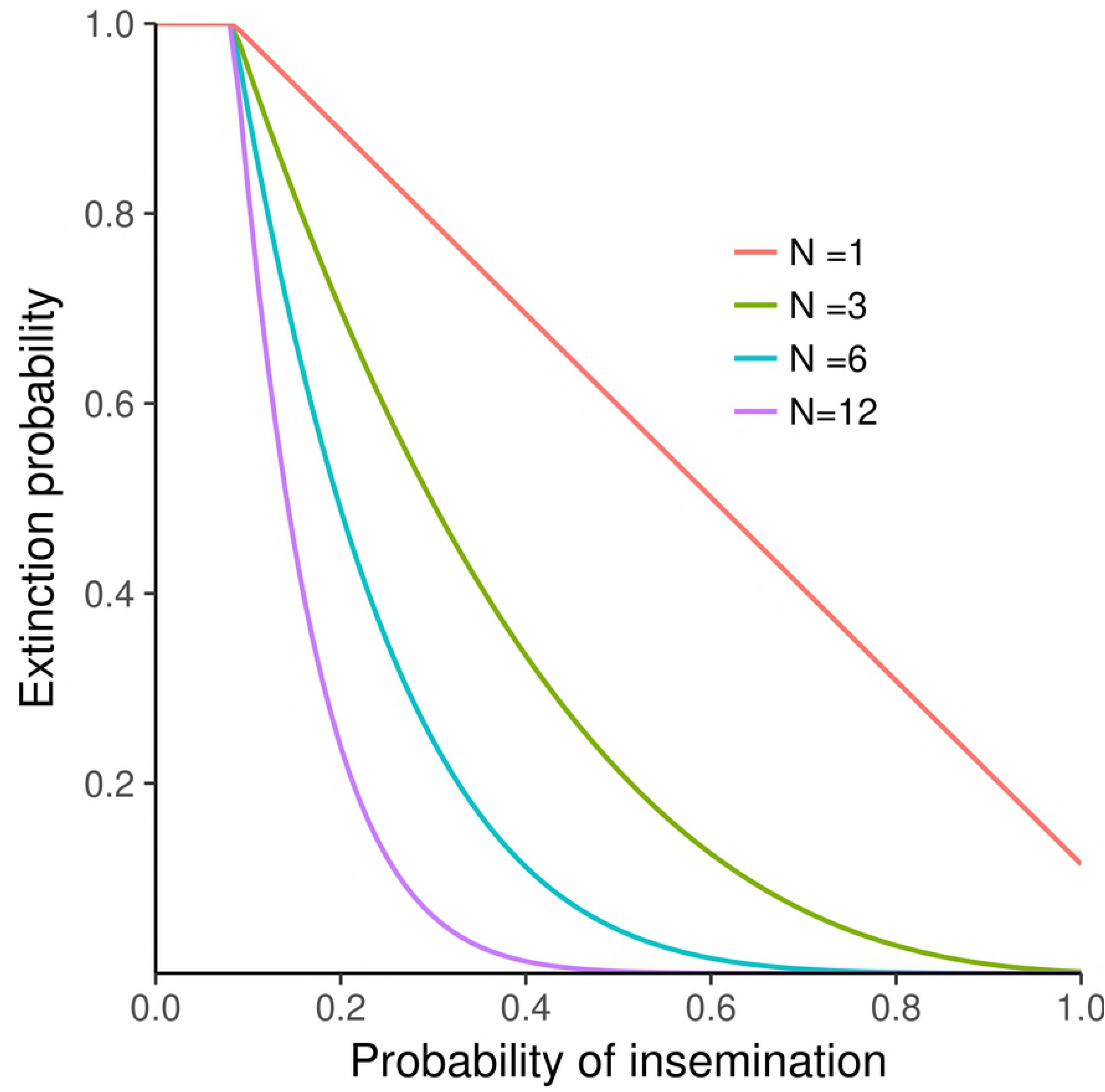
Extinction probability as a function of the probability that a female is inseminated by a fertile male, and the number of inseminated females in the pioneer population. Probability deposited pupa is female, *β* = 0.5; adult mortality rate *ψ* = 0.005 per day; pupal mortality rate *χ* = 0.005 per day; pupal duration, *P* = 27 days; time to first ovulation, *ν* = 7 days; inter-larval period *ν* = 9 days. Figures in the body of the plot show the numbers (N) of inseminated females in the pioneer population (cf [3], Fig 2B)

**Fig S5.**
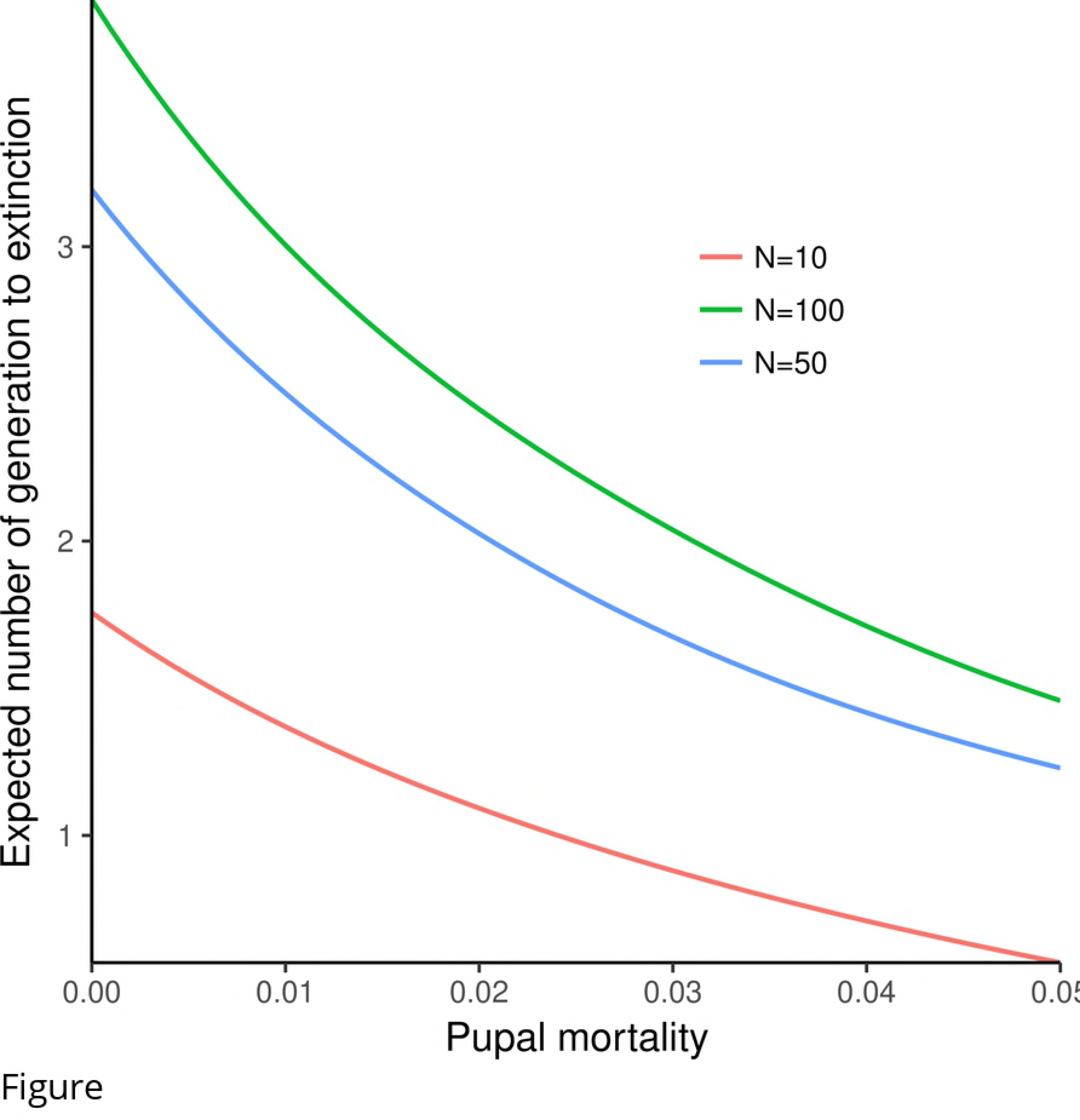
Expected number of generations to extinction as a function of pupal female mortality rate, and the number of inseminated females in the pioneer population. Input assumptions: Adult mortality rate *ψ* = 0.07 per day; probability deposited pupa is female, *β* = 0.5 probability females inseminated by a fertile male, *∊* = 1.0; pupal duration, *P* = 27 days; time to first ovulation, *ν* = 7 days; inter-larval period *τ* = 9 days. Figures in the body of the plot show the number (N) of inseminated females in the pioneer population. (cf [3], Fig 5A)

**Fig S6.**
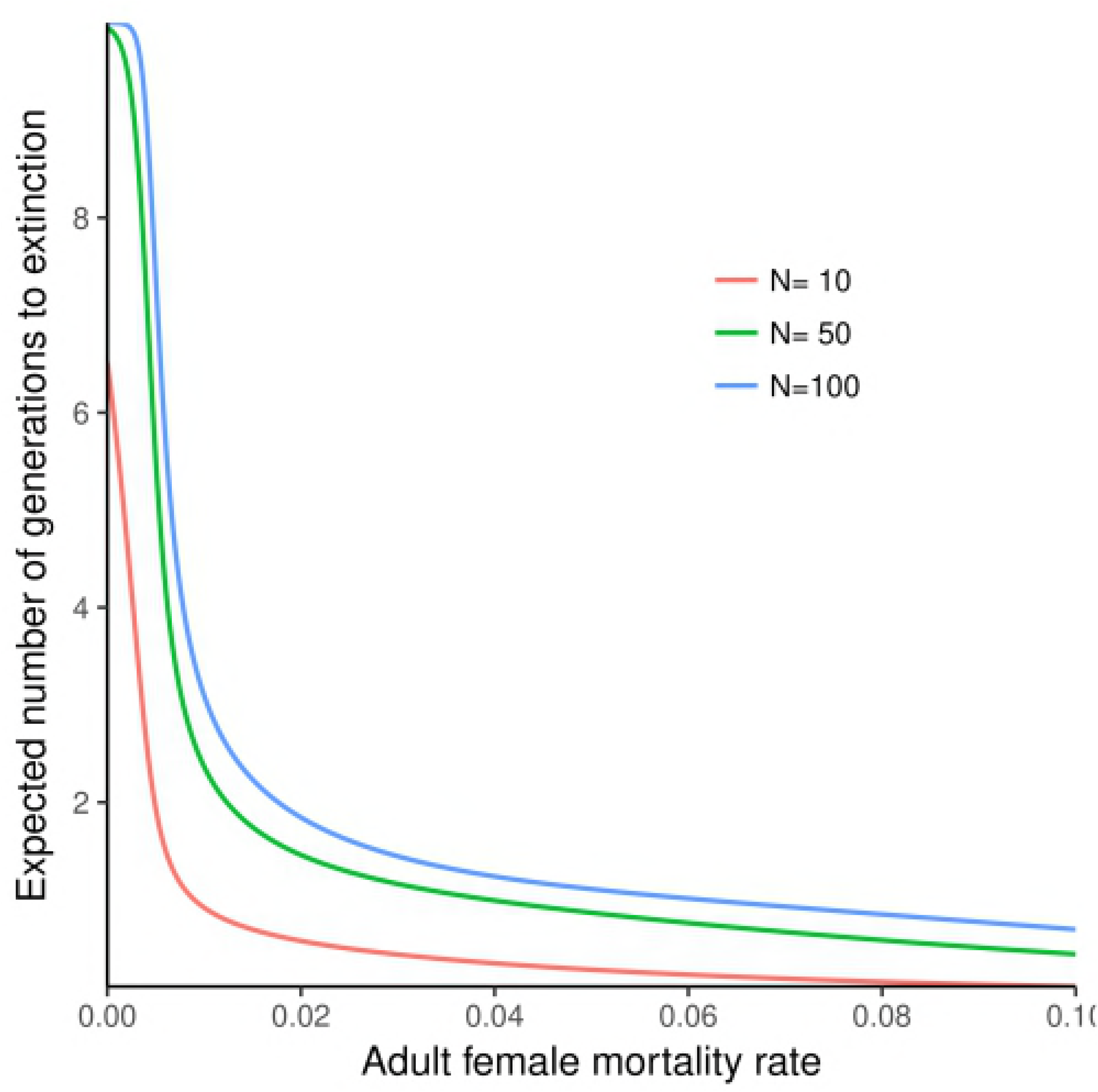
Expected time to extinction as a function of adult female mortality rate, and the number of inseminated females in the pioneer population. Input assumptions: Pupal mortality rate *ψ* = 0.005 per day; probability deposited pupa is female, *β* = 0.5; probability females inseminated by a fertile male, *∊* = 0.1; pupal duration, *P* = 27 days; time to first ovulation, *ν* = 7; inter-larval period *τ* = 9 days. Figures in the body of the plot show the number (N) of inseminated females (cf [3], Fig 5A)

**Fig S7.**
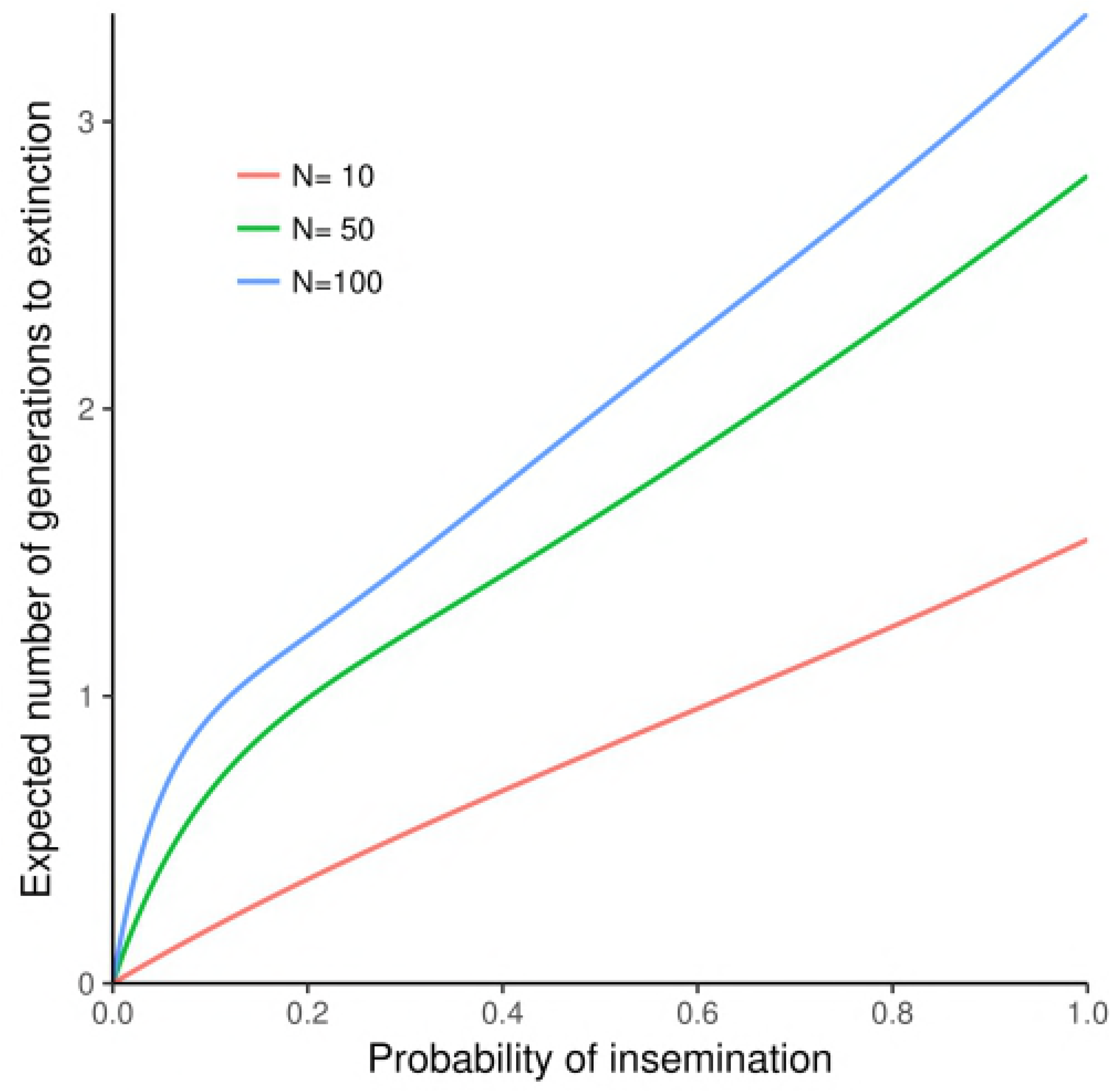
Expected time to extinction as a function of the probability that a female is inseminated by a fertile male, and the number of inseminated females in the pioneer population. Input assumptions: Adult mortality rate *ψ* = 0.07 per day; pupal mortality rate *χ* = 0.005 per day; probability deposited pupa is female, *β* = 0.5; pupal duration, *P* = 27 days; time to first ovulation, *ν* = 7 days; inter-larval period *τ* = 9 days. Figures in the body of the plot show the number (N) of inseminated females. (cf [3], Fig 5A)

## Supporting information

### SI 1: Proof of equation (7)

When *k* = 0, we obtain

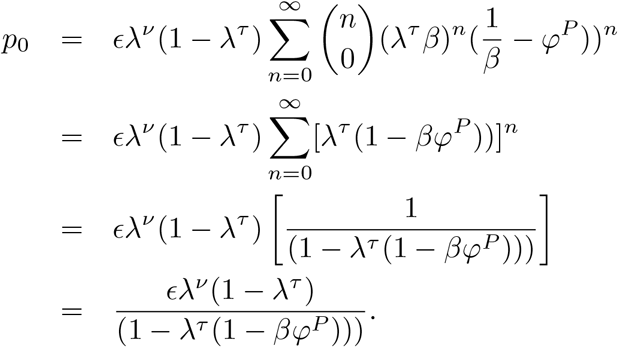

When *k* = 1, we obtain

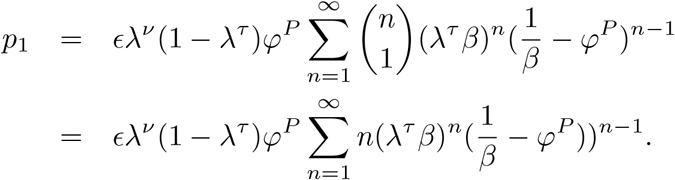

If we let 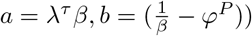 and 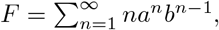, this implies that

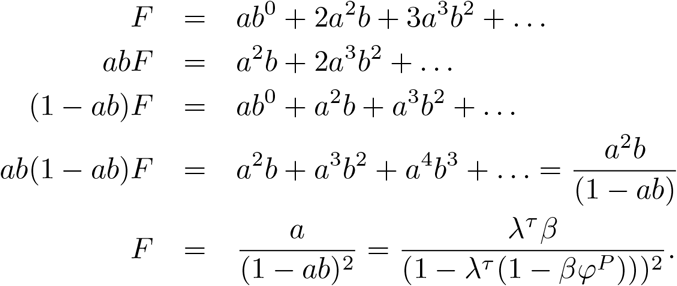

Thus, the final solution for *p*_1_ becomes

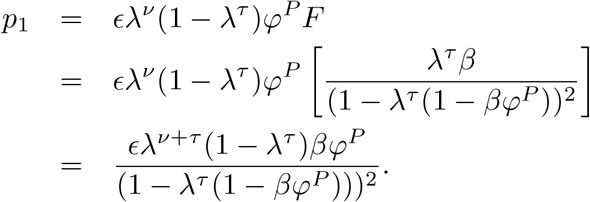

When *k* = 2, we obtain

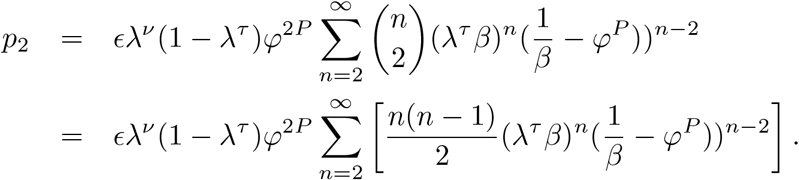

Also letting 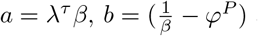 and 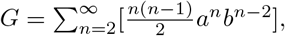, we have

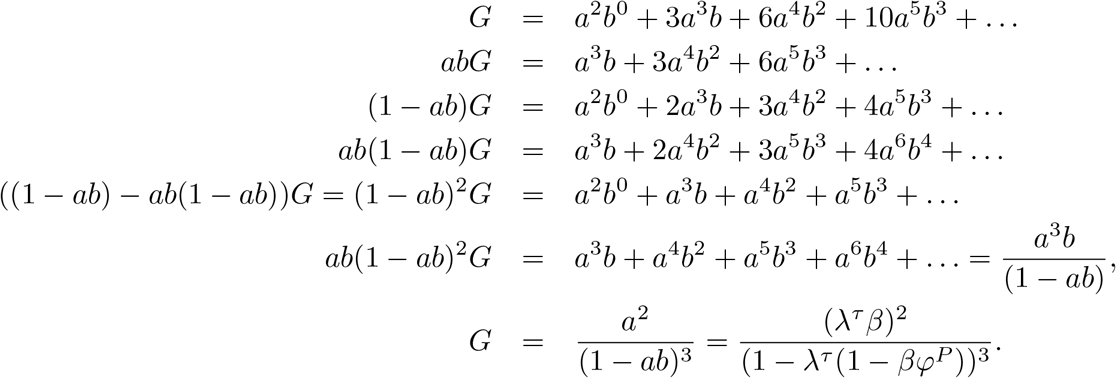

Thus, the final solution for *p*_2_ becomes

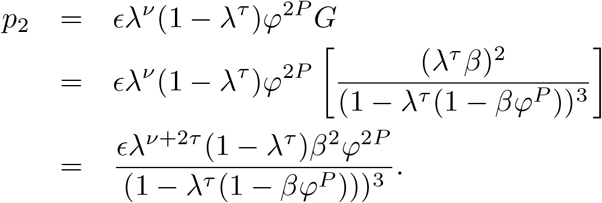

Thus, in general

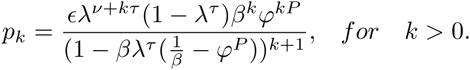

### SI 2: Proof of equations (8 and (10)

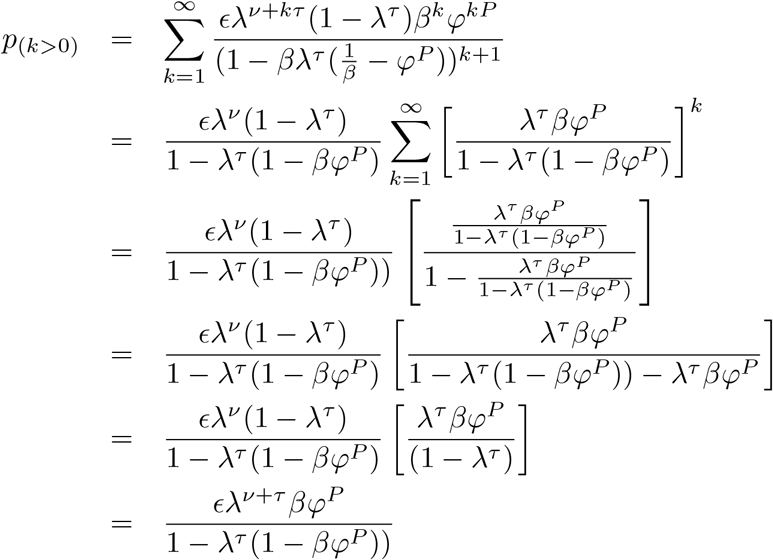

Thus, the probability that a female tsetse fly does not produce any surviving female offspring before she dies is given by:

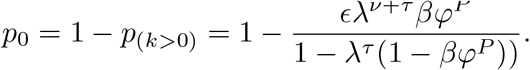

Extinction probability, *ϕ*(*θ*) is:

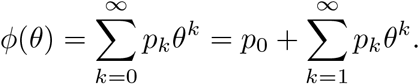

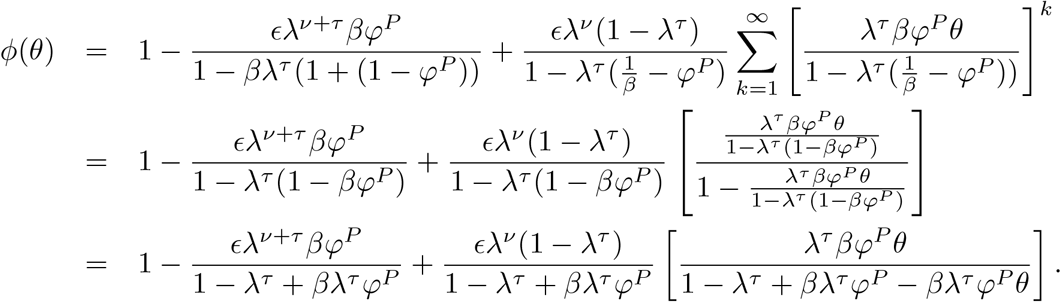

Setting *A* = 1 *− λ^τ^*, *B* = *βλ^τ^ φ^P^* and *C* = 1 *– ∊ λ^ν^*, we obtain

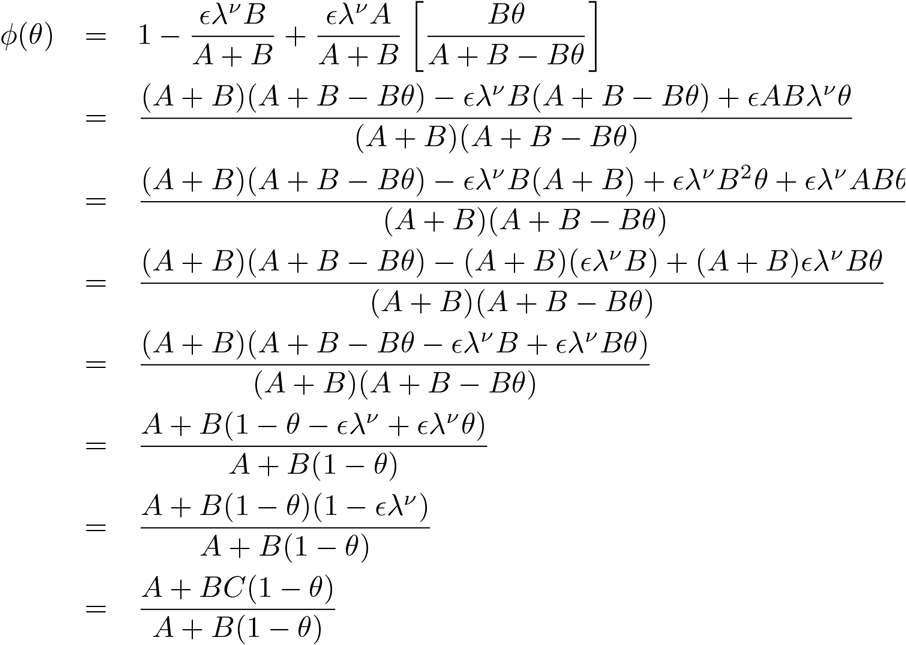

### SI 3: Proof of equations (12) and (13)

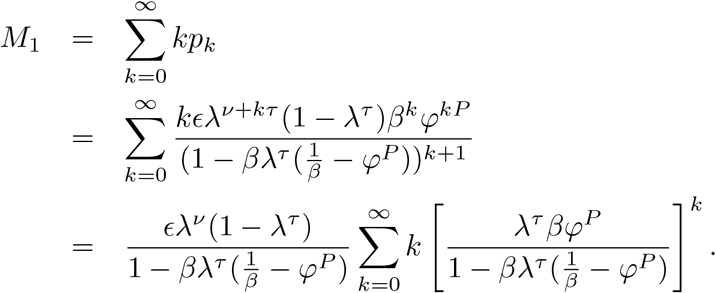

Using the sum of power series, that is 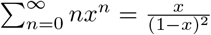 to simplify the terms not involing the summation sign, we obtain

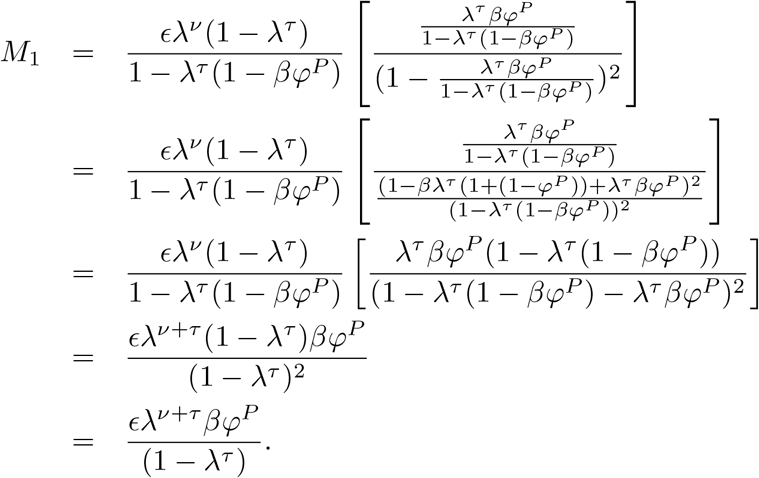

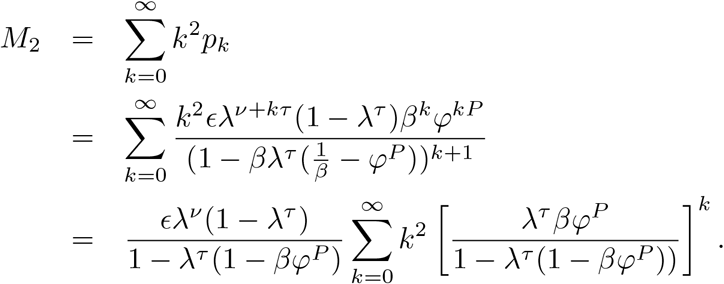

Using the sum of power series, that is 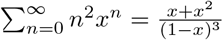 to simplify the terms not involving the summation sign, we have

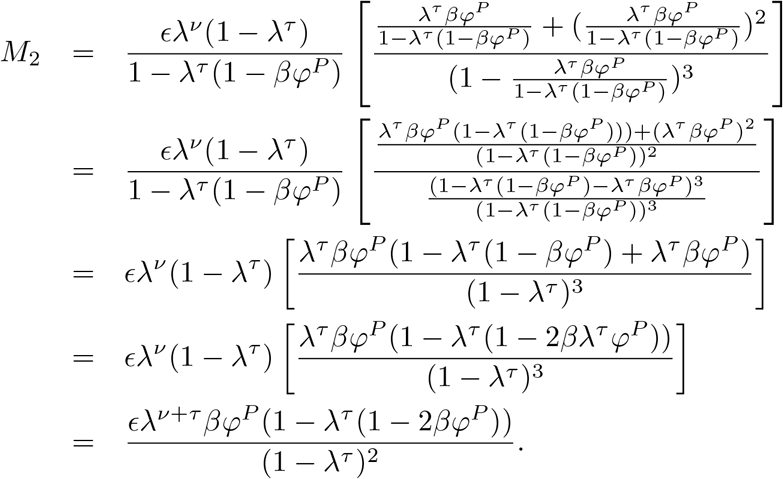

